# Detecting Novel Sequence Signals in Targeting Peptides Using Deep Learning

**DOI:** 10.1101/639203

**Authors:** J.J. Almagro Armenteros, M. Salvatore, O. Emanuelsson, O. Winther, G. von Heijne, A. Elofsson, H. Nielsen

## Abstract

In bioinformatics, machine learning methods have been used to predict features embedded in the sequences. In contrast to what is generally assumed, machine learning approaches can also provide new insights into the underlying biology. Here, we demonstrate this by presenting TargetP 2.0, a novel state of art method to identify N-terminal sorting signals, which direct proteins to the secretory pathway, mitochondria and chloroplasts or other plastids.

By examining the strongest signals from the attention layer in the network, we find that the second residue in the protein, i.e. the one following the initial methionine, has a strong influence on the classification. When subsequently examining all targeting peptides, we observe that two-thirds of chloroplast and thylakoid transit peptides have an alanine in position two, but only 20% of other plant proteins. Further highlighting the importance of the second residue, we also note that in fungi and single-celled eukaryotes, less than 30% of the targeting peptides have an amino acid that allows the removal of the N-terminal methionine compared with 60% for the proteins without targeting peptide.

TargetP 2.0 is available at http://www.cbs.dtu.dk/services/TargetP-2.0/index.php

## 1. Introduction

The localisation of proteins in the cell is a fundamental determinant of protein function. Specific sorting signals drive the subcellular localisation of proteins. These signals vary in structure, length and position between the different subcellular compartments. One of the most common types of sorting signals are the N-terminal targeting peptides. These signals are responsible for sorting proteins to the secretory pathway, mitochondria, chloroplasts (or other plastids) and compartments inside the chloroplast such as thylakoids. Signal peptides (SP) are responsible for transporting proteins to the endoplasmic reticulum to enter the secretory pathway. SPs are composed of three regions: a positively charged domain or n-region, a hydrophobic core or h-region and a segment prior to the cleavage site or c-region [1].

Mitochondrial transit peptides (mTP) are responsible for targeting proteins to the mitochondrial matrix. mTPs are usually enriched in arginine, leucine and serine. Moreover, they tend to form an amphiphilic helical structure to interact with the import receptor on the mitochondrial membrane [2]. Proteins targeted to the inner mitochondrial membrane or the inter-membrane space often have a bipartite mTP, where the second part is similar to an SP [3].

Chloroplast transit peptides (cTP) are involved in the transport of proteins to the chloroplast stroma. Most of the cTPs consist of three regions: an uncharged N-terminal region, a central region lacking acidic amino acids but enriched in serine and threonine and a C-terminal region enriched in arginine that forms an amphiphilic *β* strand [4]. Additionally, chloroplastic proteins targeted to the thylakoid lumen have a bipartite pre-sequence structure [5]. Once the cTP is cleaved and the protein enters the stroma, a luminal transit peptide (lTP) is recognised, and the protein is further transported to the thylakoid, where the lTP is cleaved. The lTP is similar to a bacterial SP, and the thylakoidal processing peptidase belongs to the family of signal peptidases [6].

As these signals direct the transport of proteins within the cell, it is crucial to be able to predict their presence in protein sequences accurately. For this reason, in the last two decades, many tools have been developed. Those adopt various machine learning algorithms including Grammatical Restrained Hidden Conditional Random Fields, N-to-1 Extreme Learning Machines, Support Vector Machines, Markov chains, profile hidden Markov models, and neural networks [7, 8, 9, 10, 11].

One of the most used methods is TargetP 1.1 [11]. TargetP uses feed-forward networks and position-weight matrices to process windows of amino acids to predict the presence of SPs, mTPs and cTPs and the positions of their cleavage sites. However, with the rise of deep learning, new types of networks such as recurrent neural networks (RNNs) have gained popularity. The main reason is their extraordinary ability to work with sequence data and model long-range relationships between inputs in the sequence.

RNNs sequentially process sequences of any length, being able to retain information from previous positions in the sequence. Several methods have taken advantage of this type of network to try to better predict signal and transit peptides [12, 13]. These methods make use of bidirectional RNNs (BiRNN), which are two RNNs, one processing the sequence forwards and another processing the sequence backwards. With this construction the context around each amino acid is modelled, as the forward RNN processes all the amino acids from the N-terminus up to one position and the backward RNN processes all the amino acids from the C-terminus up to the same position.

However, regular RNNs, so-called Elman networks, are challenging to train (the so-called exploding/vanishing gradient problem) and often fail to capture dependencies far apart in the sequence [14] Therefore, the ability of the network to hold information from multiple steps back is reduced. A variant of the RNN cell, the Long Short-Term Memory (LSTM), solves this problem by a construction akin to a computer memory cell that holds information for multiple steps. This type of RNN cell together with BiRNN have been successfully applied to the prediction of SPs and mTPs [15, 16]. Today, new methods such as DeepLoc [17] uses bidirectional LSTM to predict the localisation of proteins to a broader range of compartments. DeepLoc accurately predicts the localisation of proteins but not the presence of the N-terminal sorting signals and the position of the cleavage sites. Starting from this architecture, we decided to develop TargetP 2.0 using bidirectional LSTM and a multi-attention mechanism. Using the multi-attention mechanism the network can predict both the type of peptide and the position of the cleavage site by focusing on particular regions of the sequence.

Moreover, we assemble a new protein dataset that we use to train TargetP 2.0. TargetP 2.0 can jointly predict the presence of signal peptides, mitochondrial, chloroplast and thylakoid transit peptides, and the corresponding cleavage site positions.

When analysing the attention layer from the final version of the network, it became apparent that most information was retrieved from two distinct positions in most sequences. One of these was, as expected, localised close to the cleavage site. However, an equally important signal from position two in the sequences was also found. Next, we examined the amino acid frequencies in the second position, after the first methionine, of all proteins. To our surprise, very distinct patterns emerged. In chloroplasts and plastids, the second residue was frequently an alanine, while in all targeting peptides in fungi and unicellular eukaryotes amino acids that allow cleavage of the methionine are rare, see Figure 1.

**Figure 1:**
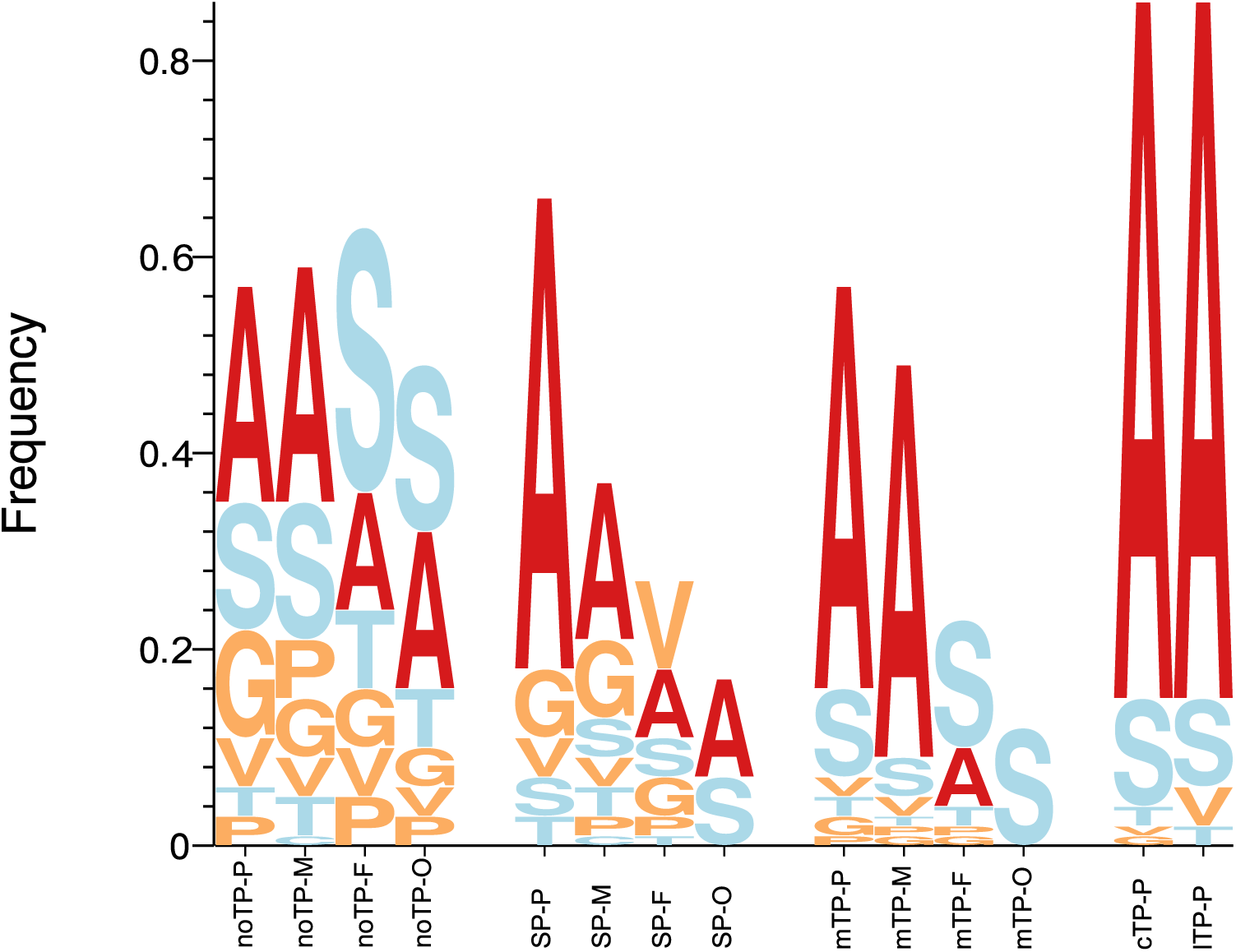
This figure depicts the frequencies of the second residue in proteins with different targeting peptides. The proteins are divided into their respective type of targeting peptide (Signal Peptides (SP), mitochondrial targeting peptides (mTP), chloroplast targeting peptides (cTP), thylakoid targeting peptides (lTP) and proteins without a targeting peptide (noTP). Further, the proteins were divided into their kingdom: Viridiplantae (P), Metazoa (M), Fungi (F) and other Eukaryotic organisms (O) sequences. Inspired by sequence LOGOs the height of each letter corresponds to the frequency of that amino acid. Only the frequencies for the short side-chained amino acids that allow the cleavage of the N-terminal methionine are shown.

## 2. Material and methods

### 2.1. Dataset

The protein data used to train TargetP 2.0 were extracted from the UniProt database, release 2018 04 [18]. The negative dataset consists of proteins without either signal or transit peptides from the nucleus, cytoplasm and plasma membrane (without signal peptides) and with experimental annotation (ECO:0000269) of their subcellular localisation. The positive set contained secreted, mitochondrial, chloroplastic and luminal proteins with experimental annotation of their signal or transit peptide. The final set consists of 9537 (noTP) proteins without targeting peptides, 2697 with SPs; 499 mTPs, 227 cTPs and 45 lTPs, see Table 1. Note that although a thylakoid targeting signal, as described in the introduction, consists of a cTP followed by an SP-like lTP, the first cleavage site (for the stromal processing peptidase) is almost never annotated in UniProt. We are therefore not able to predict this cleavage site for thylakoid proteins, only the second cleavage by thylakoidal processing peptidase will be predicted. Hereafter, “lTP” will refer to the entire thylakoid targeting signal. The dataset was further divided into four groups representing the eukaryotic kingdoms Viridiplantae, Metazoa, and Fungi and a group of other eukaryotes.

**Table 1:** Performance of the predictors considering only the identification of the targeting peptides. The table shows the performance in the test set yield by each predictor for Mitochondria (mTP), Chloroplast (cTP), Thylakoid (lTP), Signal Peptide (SP), and Other (noTP), in terms of F1 score, Matthews correlation coefficient (MCC), Precision (Prec) and Recall (Rec).

|  | **Tool** | **Loc** | **Proteins** | **Prec** | **Rec** | **F1-Score** | **MCC** |
| --- | --- | --- | --- | --- | --- | --- | --- |
|  | **TargetP 2.0** | **SP** | 2697 | 0.97 | 0.98 | 0.98 | 0.97 |
|  | **TargetP 1.1** | **SP** | 2697 | 0.86 | 0.97 | 0.91 | 0.89 |
|  | **DeepLoc** | **SP** | 2697 | 0.90 | 0.84 | 0.87 | 0.84 |
|  | **PredSL** | **SP** | 2697 | 0.69 | 0.90 | 0.78 | 0.73 |
|  | **Predotar** | **SP** | 2697 | 0.92 | 0.92 | 0.92 | 0.90 |
|  | **MLP-20** | **SP** | 2697 | 0.93 | 0.93 | 0.93 | 0.91 |
|  | **TargetP 2.0** | **mTP** | 499 | 0.87 | 0.85 | 0.86 | 0.86 |
|  | **TargetP 1.1** | **mTP** | 499 | 0.32 | 0.90 | 0.48 | 0.51 |
|  | **DeepLoc** | **mTP** | 499 | 0.73 | 0.97 | 0.83 | 0.83 |
|  | **PredSL** | **mTP** | 499 | 0.18 | 0.93 | 0.31 | 0.37 |
|  | **Predotar** | **mTP** | 499 | 0.71 | 0.74 | 0.73 | 0.72 |
|  | **TPPred3** | **mTP** | 499 | 0.69 | 0.68 | 0.68 | 0.67 |
|  | **Mitofates** | **mTP** | 499 | 0.70 | 0.92 | 0.80 | 0.80 |
|  | **MLP-20** | **mTP** | 499 | 0.69 | 0.58 | 0.63 | 0.62 |
|  | **TargetP 2.0** | **cTP** | 227 | 0.90 | 0.86 | 0.88 | 0.88 |
|  | **TargetP 1.1** | **cTP** | 227 | 0.39 | 0.88 | 0.54 | 0.58 |
|  | **DeepLoc** | **cTP** | 227 | 0.70 | 0.94 | 0.80 | 0.80 |
|  | **PredSL** | **cTP** | 227 | 0.16 | 0.78 | 0.27 | 0.34 |
|  | **Predotar** | **cTP** | 227 | 0.51 | 0.76 | 0.61 | 0.61 |
|  | **TPPred3** | **cTP** | 227 | 0.76 | 0.64 | 0.69 | 0.69 |
|  | **MLP-20** | **cTP** | 227 | 0.51 | 0.37 | 0.43 | 0.40 |
|  | **TargetP 2.0** | **ITP** | 45 | 0.75 | 0.75 | 0.75 | 0.75 |
|  | **PredSL** | **ITP** | 45 | 0.46 | 0.71 | 0.56 | 0.57 |
|  | **MLP-20** | **ITP** | 45 | 0.10 | 0.02 | 0.04 | 0.05 |
|  | **TargetP 2.0** | **noTP** | 9537 | 0.98 | 0.98 | 0.98 | 0.95 |
|  | **TargetP 1.1** | **noTP** | 9537 | 0.99 | 0.84 | 0.91 | 0.75 |
|  | **DeepLoc** | **noTP** | 9537 | 0.95 | 0.95 | 0.95 | 0.83 |
|  | **PredSL** | **noTP** | 9537 | 0.99 | 0.60 | 0.75 | 0.52 |
|  | **Predotar** | **noTP** | 9537 | 0.96 | 0.95 | 0.95 | 0.84 |
|  | **TPPred3** | **noTP** | 9537 | 0.76 | 0.98 | 0.86 | 0.29 |
|  | **Mitofates** | **noTP** | 9537 | 0.75 | 0.98 | 0.85 | 0.25 |
|  | **MLP-20** | **noTP** | 9537 | 0.95 | 0.97 | 0.96 | 0.85 |

PSI-CD-HIT [19] was used to cluster the first 200 residues of each protein with 20% of identity or 10^−6^ E-value using BLAST and alignment coverage of at least 80% of the shorter sequence. We performed a stringent homology partitioning to get a realistic assessment of generalisation performance. Each cluster of homologous proteins was assigned to one of five cross validation groups to ensure that similar proteins were not mixed between the different datasets.

### 2.2. The TargetP 2.0 algorithm

The TargetP 2.0 model is described in Figure 2. The model consists of two key components, a bidirectional recurrent neural network with LSTM cells and a multi-attention mechanism [20] to predict both the type of peptide and the position of the cleavage site.

**Figure 2:**
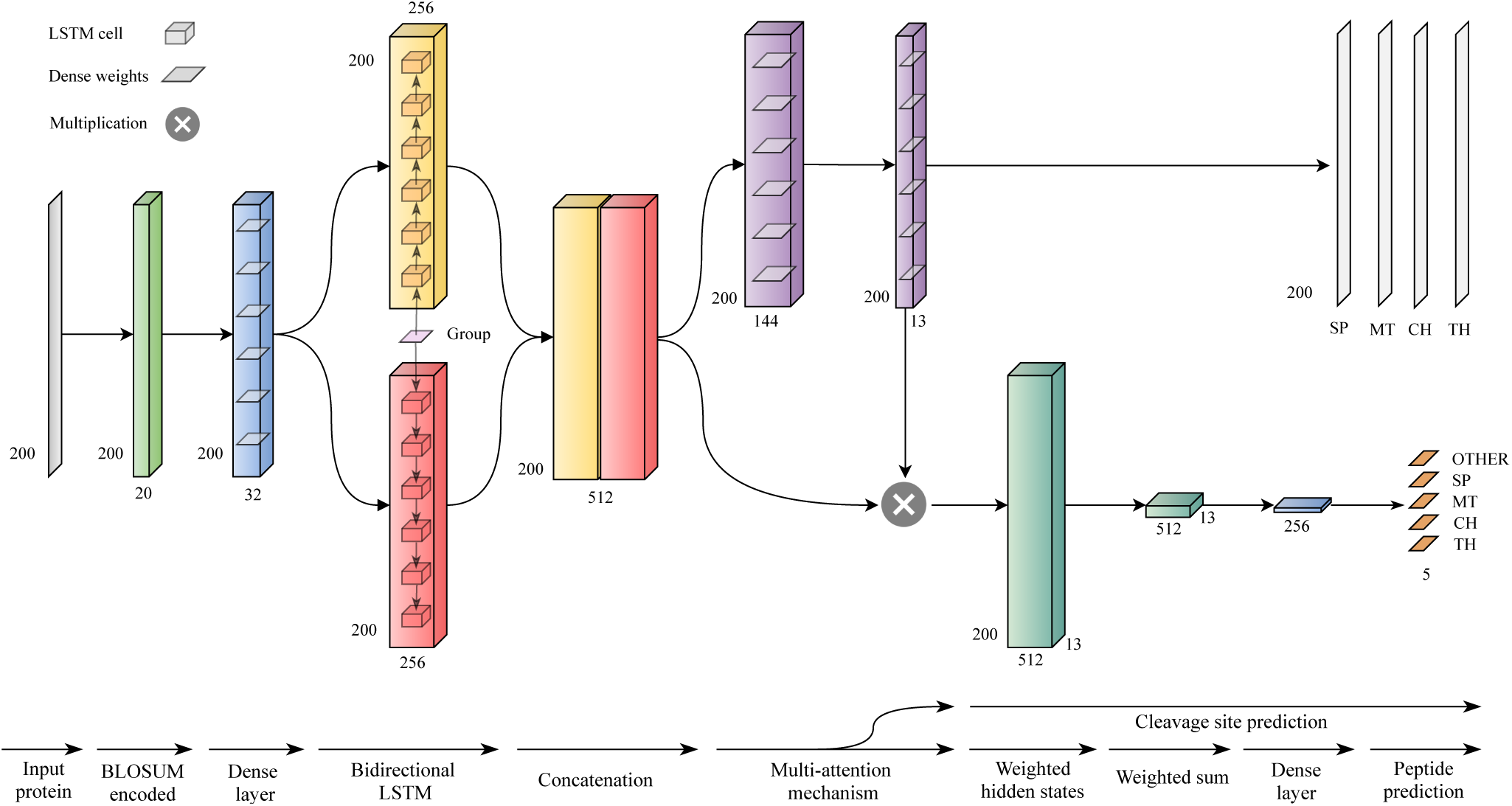
The TargetP 2.0 architecture.

The input to this model is the first 200 amino acids of a protein. This threshold was chosen based on the maximum length of known transit peptides, which is 162 amino acids [21]. The amino acids in the protein are encoded using BLOSUM62 substitution matrices.

We first describe the model at a high level and give more details on each of the layers below: The first layer of the model is a fully connected layer to perform a feature transformation of each amino acid input feature with 32 hidden units. The following layer is the bidirectional LSTM (BiLSTM) with 256 hidden units in both forward and backward direction. The first hidden state to the BiLSTM is a vector containing the group information, which denotes whether the protein is plant or non-plant. The 512-dimensional concatenated output from the BiLSTM is then used to calculate the multi-attention matrix similarly to those applied in machine translation [22, 23]. The attention size is 144 units and the number of outputs from the attention matrix of size 13. Out of these 13 attention vectors, 4 were used to predict the different cleavage site positions for SP, mTP, cTP and lTP. The attention matrix is further utilised to encode the whole sequence into a context matrix. This context matrix of size 512×13 is processed by a fully connected layer with 256 units, to summarise it into a vector. Finally, this is fed to the output layer with 5 hidden units and softmax activation.

We train a model that learns to predict the type of peptide and the position of the corresponding cleavage site (*y, y*′) = *f*_*θ*_(*X*) where *y* is the predicted type of peptide, *y*′ the predicted cleavage site position, *f* the model, *θ* the learnable parameters and *X* the protein sequence. Here, *y* is a vector of size equal to the number of classes *C*, five in this case, and *y*′ is a vector of size equal to the length of the sequence *L*, which can be up to 200. The *θ* parameters are optimised using an extension of stochastic gradient descent, ADAM with cross-entropy loss for both types of peptide and cleavage site prediction. Both losses were then averaged. The only regularisation technique used was dropout between the different layers.

The network has three main types of layers: fully connected, RNN with LSTM cell and multi-attention layer. The first fully connected layer *c* applies a feature transformation:

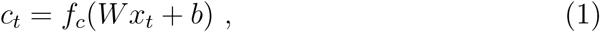

where *x*_*t*_ is an amino acid at position *t* in the sequence and *W* and *b* the learnable weights and biases. The first layer is followed by a bidirectional RNN that utilises an LSTM cell to capture the context around each amino acid in the sequence. The RNN applies the same set of weights to each position *t*

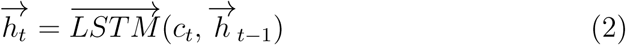

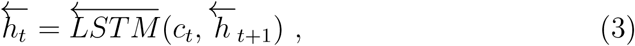

where 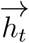 and 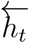 are the hidden states of the RNN at position *t* for the forward and backward direction respectively. The hidden states are concatenated into 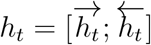.

The last part of the network is a multi-attention mechanism. Here we calculate multiple attention vectors *A* from the LSTM hidden states, instead of just one single attention vector *a*. The attention matrix is then used to create multiple fixed sized representations of the input sequence, with a different focus on the relevant parts of the sequences. The attention matrix is calculated as follows:

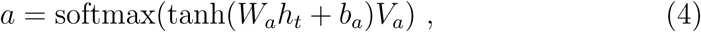

where *W*_*a*_ and *V*_*a*_ are weight matrices and *b*_*a*_ is the bias of the attention function. The advantage of having multiple attention vectors is that some of them can be used to predict the position of the cleavage site, as they are vectors of size equal to the sequence length *L* summing to one. Therefore, 4 out of the 13 attention vectors that the model uses are employed in the prediction of the SP, mTP, cTP and lTP cleavage site (cs):

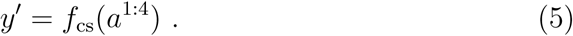

To encode the sequence of hidden states *H* = [*h*_1_, …, *h*_*L*_] into a fixed sized matrix, the hidden states are multiplied by the attention matrix and summed up:

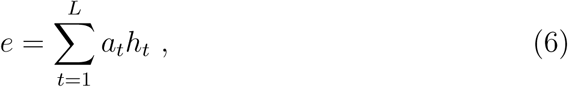

where *e* matrix is the encoded representation of the protein sequence. *e* holds a total of 13 different representations of the protein sequences; therefore, it is needed to summarise this matrix into a vector. This is done by a final feed-forward layer, which converts *E* into a representation vector *e*. This is then used to calculate the output layer of the network, to predict the type of peptide (p) *y*

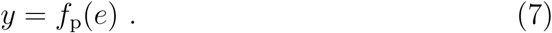

Both outputs from the network *y* and *y*′ are trained together. The exception is for proteins belonging to the negative set, i.e. proteins without targeting peptides that lack a cleavage site and therefore there is no error to back-propagate.

The model was trained and optimised using five-fold nested cross-validation. The four inner subsets were used to train the model, where three are used for training and one for validation to identify the best set of hyper-parameters. After optimisation, the fifth set, that was kept out of the optimisation, was used to assess the test set performance. This procedure was repeated using all five subsets as the test set. The advantage of this approach is that we obtain an unbiased test set performance on the whole dataset at the expense of having to train 5×4=20 models.

Different hyper-parameters were tested to find the best model such as the number of hidden units for the LSTM, attention and fully connected layers, number of attention vectors, the learning rate and dropout rate. We also experimented with a convolutional neural network (CNN) as the initial layer, but the best results were achieved using a filter size of 1, which is equivalent to a fully connected layer along the feature dimension.

### 2.3. Related tools

The tools included in the analysis adopt different machine learning algorithms intending to classify from one to many N-terminal sorting signals and the cleavage site position. Most of the tools contain modules both for plant and non-plant proteins.

**TargetP 1.1** [11, 24] classifies proteins into 4 different groups (signal peptide, mitochondrial transit peptide, chloroplastic transit peptide and other) using two layers of feed-forward neural networks and detects the cleavage sites using a variety of methods including postion-weight matrices for the mTPs.

**TPPred3** [7] is a combination of a Grammatical Restrained Hidden Conditional Random Field, N-to-1 Extreme Learning Machines and Support Vector Machines. It detects transit peptides, classifying them as mitochondrial or chloroplastic and localising their cleavage sites.

**Mitofates** [8] combines amino acid composition and physico-chemical properties with positively charged amphiphilicity, pre-sequence motifs, and position weight matrices as input to a standard support vector machine classifier for modelling the mitochondrial pre-sequence and its cleavage site.

**PredSL** [9] uses neural networks, Markov chains, profile hidden Markov models, and scoring matrices to classify proteins from the N-terminal amino acid sequence into five groups: chloroplast, thylakoid, mitochondrion, secretory pathway, and other.

For comparison, we also choose to include two methods that do identify the subcellular localisation of proteins but do not predict the cleavage site of the targeting peptides.

**Predotar** [10] is a three-layer feed-forward neural network-based approachable to classify proteins in 4 different classes: signal peptide, mitochondrial transit peptide, chloroplast transit peptide and other.

**DeepLoc** [17] uses a deep learning architecture very similar to what we have used in this study to predict the subcellular localisation of proteins.

**MLP-XX** is a simple multi-layer perceptron that we tested for comparison. MLP-XX consists of a one layer feed forward neural network where using one hot encoding of the first XX amino acids as input (up to 20). It used the same cross-validation as TargetP 2.0. We examined the inclusion of different numbers of N-terminal residues, and the average F1-score increased from 0.77 when using five residues to 0.93 when using twenty, see supplementary table S1. For comparison we include MLP-20 in the results.

#### 2.3.1. Evaluation of the performance

We use several performance measures to obtain a uniform evaluation of the prediction. For the performance of sorting signals, we use the F1 score that may count as a harmonic average of the precision and recall. We also computed the Matthews Correlation Coefficient (MCC) for each class, to have a much more balanced evaluation of the prediction [25]. On the other hand, we use precision and recall for the combined performance of sorting signals and cleavage site. All these measurements were expressed in terms of “tp” = true positive, “tn” = true negative, “fp” = false positive,”fn” = false negative.

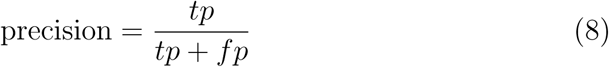

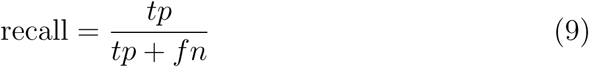

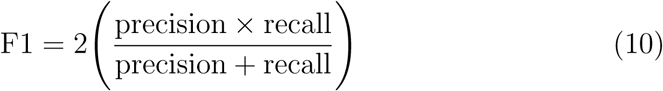

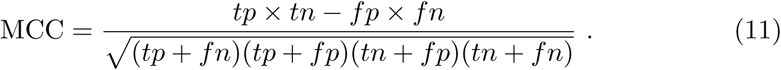

### 2.4. Additional analysis

In several figures standard or variations of sequence LOGOs are shown. These were generated using the Seq2Logo program [26]. In addition to standard sequence LOGOs calculated from multiple sequence alignments LOGOs representing the frequency of amino acids in position two (and not the entropy) and LOGOs representing the strength of the attention layer output were generated.

Secondary structure preferences for the different targeting peptides were calculated from the scale from [27]. The Log2 of the average preference was plotted for each residue in the different targeting peptides.

## 3. Results and Discussion

Here, we have developed a deep learning model to predict targeting peptides described in Figure 2.

First, we compare TargetP 2.0 with state-of-the-art predictors on a set of proteins with experimentally verified targeting peptides.

### 3.1. TargetP 2.0 improved identification of targeting peptides

In Table 1 it can be seen that TargetP 2.0 is better than all the competitors at the identification of targeting peptides in accuracy and correlation coefficients. From the ROC curves in Figure 3, it is clear that TargetP 2.0 performs better than the alternative methods for identification of all four targeting peptides. It can also be noted that the identification of signal peptides is more reliable than the identification of transit peptides. TargetP 2.0 predicts approximately 97% of the SPs correctly compared to less than 90% for other targeting peptides, see Table 1. For non-plant proteins the most common confusion is between mTPs and nonTPs, see Table S2.

**Figure 3:**
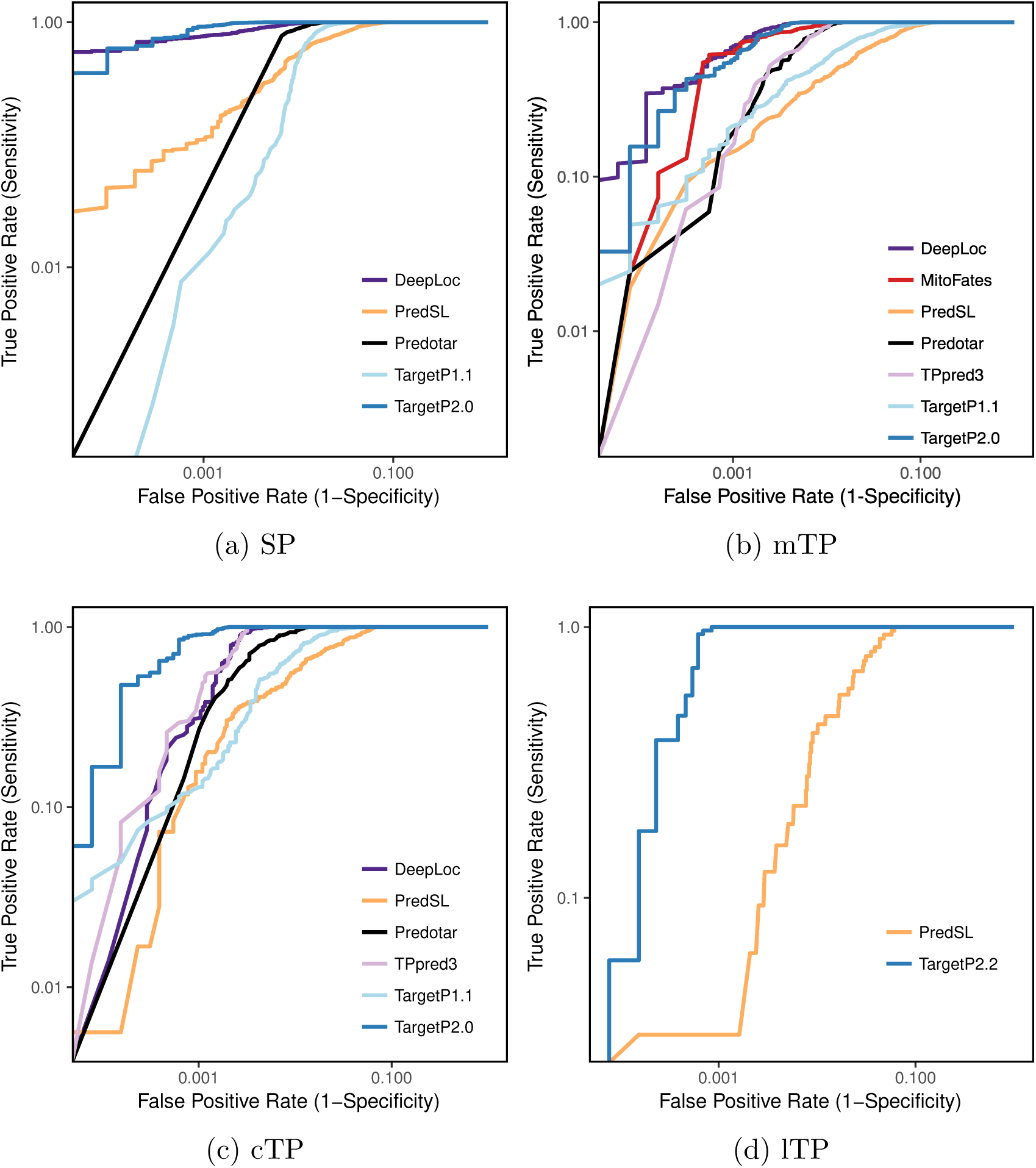
ROC curves for identification of signal peptides, mitochondrial-, chloroplast- and thylakoid targeting peptides.

The poor discrimination between mTP and cTP of TargetP 1.1 and other older methods has been significantly improved in TargetP 2.0. The number of correctly predicted peptides increased from about 50% to 90%. The only other method that shows a similar performance is DeepLoc, which is based on a similar methodology and training set but cannot predict cleavage sites. TargetP 2.0 also performs significantly better at the identification of cTPs and lTPs than PredSL [9]; the only other method that can identify lTPs. However, still 11 out of 45 lTPs are classified as cTPs, see Table S3.

It can be seen that a very simple method that only considers the 20 N-terminal amino acids, MLP-20, performs on par with previous methods when it comes to mTPs and SPs, but slightly worse than Predotar for cTPs. Even when only using ten residues, MLP-10 performs better than PredSL for all categories except lTPs, see Table S1.

A more detailed analysis at the kingdom level for TargetP 2.0 can be found in Table S4. Here, we can see that the prediction accuracy is slightly lower in Fungi than in the other kingdoms. One possible explanation could be that the GC content in the Fungi genomes is lower than for the other genomes. The low GC content affects the amino acid frequencies, making alanine less frequent [28].

Since the chloroplast is not the only type of plastid, we finally tested the ability of TargetP 2.0 to predict proteins of amyloplasts and chromoplasts, which differ from chloroplasts primarily through their pigments. UniProt provides transit peptide annotation for 10 amyloplast and 32 chromoplast proteins. TargetP 2.0 predicts 9 out of 10 amyloplast and 26 out of 32 chromoplast proteins to have a cTP, achieving a similar performance for these plastid proteins.

### 3.2. TargetP 2.0 improves the prediction of cleavage sites in cTPs and lTPs

We tested the cleavage site prediction ability on the test set and only for the correctly predicted proteins. The cleavage site prediction is best for SPs, with a recall (accuracy) of 83% on the test set both for TargetP 1.1 and TargetP 2.0, see Figure 4 and Table S5.

**Figure 4:**
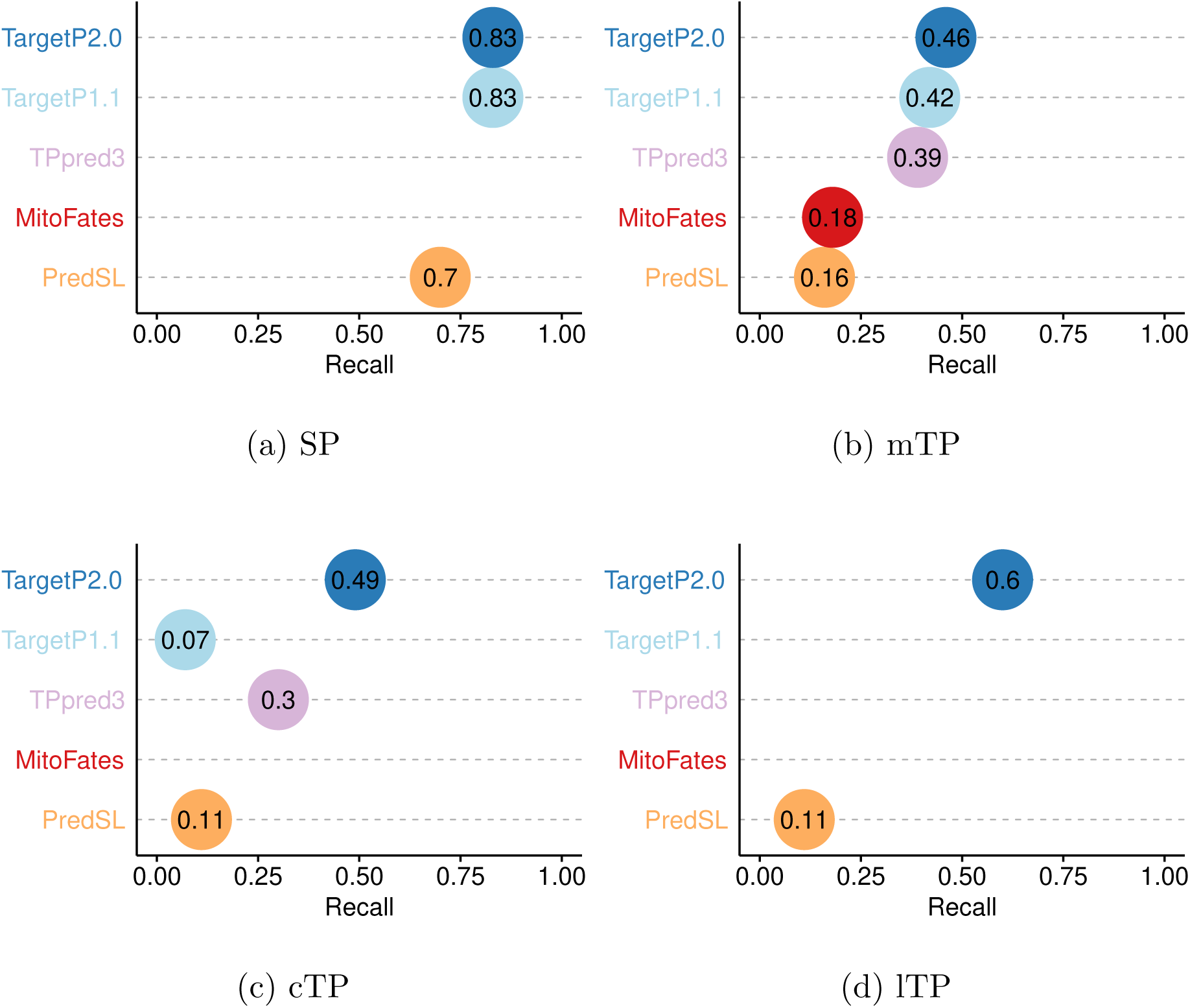
Recall (or accuracy) for the cleavage site prediction in SPs, mTPs, cTPs and lTPs by the different prediction methods. Note that not all methods can predict all types of targeting peptides.

In mTP and cTP cleavage site prediction is more difficult with a recall of 46% and 49% by TargetP 2.0, respectively. However, this is a clear improvement over TargetP 1.1 and all other methods for cTPs, and a slight improvement for mTPs.

TargetP 2.0 cleavage site predictions of the lTP is a new feature. Given the small number of peptides in the database, the recall of 60% (27 correctly identified lTP cleavage sites) is better than expected and a significant improvement over the only other method that can predict lTPs, PredSL [9], which only identifies 5 (11%) cleavage sites correctly.

If we allow for up to 5 residues shifts of the prediction of cleavage sites about two-thirds of the cleavage sites in cTPs, mTPs and lTPs can be identified correctly, see Table S5.

### 3.3. Comparison with UniProt annotations

TargetP 2.0 provides a possibility for fast and accurate annotation of entire or incomplete proteomes in a few hours, as it takes on average only 0.20 seconds to run a single protein on a dedicated 8-core machine. We annotated several eukaryotic proteomes for a total of 288964 proteins from six Metazoa (*Caenorhabditis elegans, Drosophila melanogaster, Danio rerio, Homo sapiens, Mus musculus* and *Xenopus tropicalis*), five Viridiplantae (*Arabidopsis thaliana, Brachypodium distachyon, Oryza sativa, Solanum lycopersicum* and *Vitis vinifera*) and two Fungi (*Saccharpmyces cerevisiae* and *Schizosaccharomyces pombe*) proteomes. All predictions are available from the accompanying website. We examined the possibility to modify the number of annotated proteins using the confusion matrix of TargetP 2.0 as proposed before [29]. The number of peptides in each class changed with less than 3% for all categories except the lTPs that were underpredicted by 25%, see Table S6. This indicates that our estimates of the number of targeting peptides of each type should be rather accurate except for lTPs.

In Table S7 a comparison of the annotations from TargetP 2.0 and UniProt is presented. For the best-annotated proteomes, *H. sapiens, M. musculus, S. cerevisiae* and *A. thaliana*, the agreement between UniProt and TargetP 2.0 predictions is about 80% for the organelles and over 90% for signal peptides. The high agreement for SPs is quite likely due to UniProt applying SignalP [30] for its annotation of SPs, and it was trained on a similar dataset as used here. For the other proteomes, the agreement is substantially worse, except for SPs, indicating that the transit peptide annotation in UniProt is far less complete than the SP annotation and that applying TargetP 2.0 would significantly improve the annotation.

A few interesting differences can be observed, that might have biological relevance. TargetP 2.0 predicts about twice as many mitochondrial proteins in plant proteomes compared to metazoan proteomes. Even in *A. thaliana* only half of these proteins are annotated in UniProt as mitochondrial. In agreement with the UniProt annotations fungi seem to have fewer mitochondrial proteins than other eukaryotes. The number of predicted chloroplast proteins varies significantly between the proteomes, from 1125 in the grape proteome to 2049 in the rice proteome. However, the rice proteome is also almost 50% larger than the grape proteome, possibly explaining the difference.

### 3.4. Identification of the strongest contributing sequence factors

Above we show that by using a deep learning architecture it is possible to improve the prediction of targeting peptides. Next, we wanted to examine if it is possible to extract which biological features contribute to improved performance.

To analyse which features the deep learning model learned we focused on the maximum outputs from the attention layer, see Figure S1. It is clear that for most proteins with targeting peptides there are two positions with strong signals, one very close to the N-terminus (at position 2) and one later corresponding to a position close to the cleavage site. These positions were analysed in more detail, by aligning all the proteins either starting from the predicted cleavage site, Figure 5, or from the N-terminus, Figure 6.

**Figure 5:**
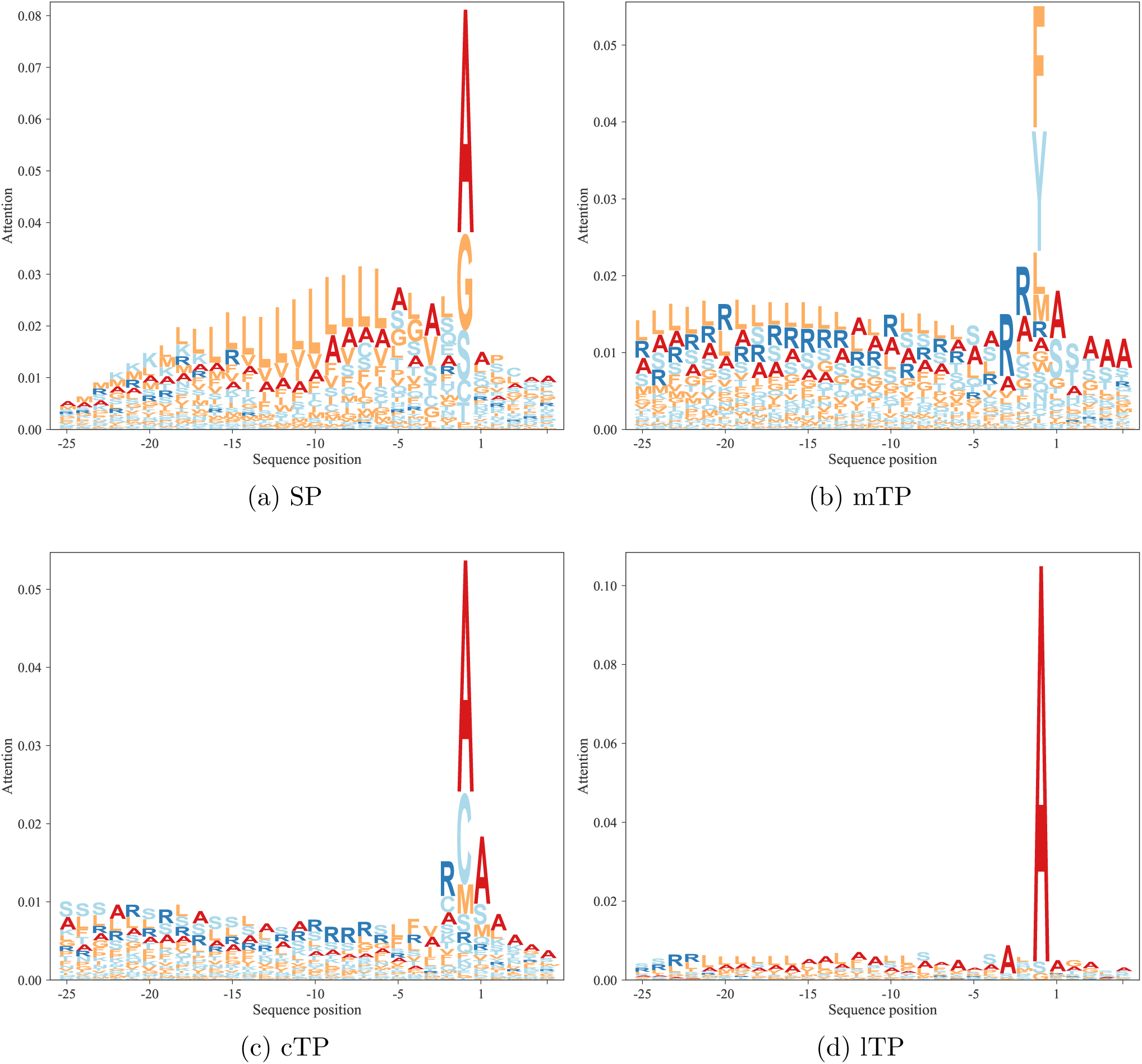
Attention layer LOGOs showing the impact strength of the attention layer and the frequency of amino acids. All sequences are aligned at the predicted cleavage site.

**Figure 6:**
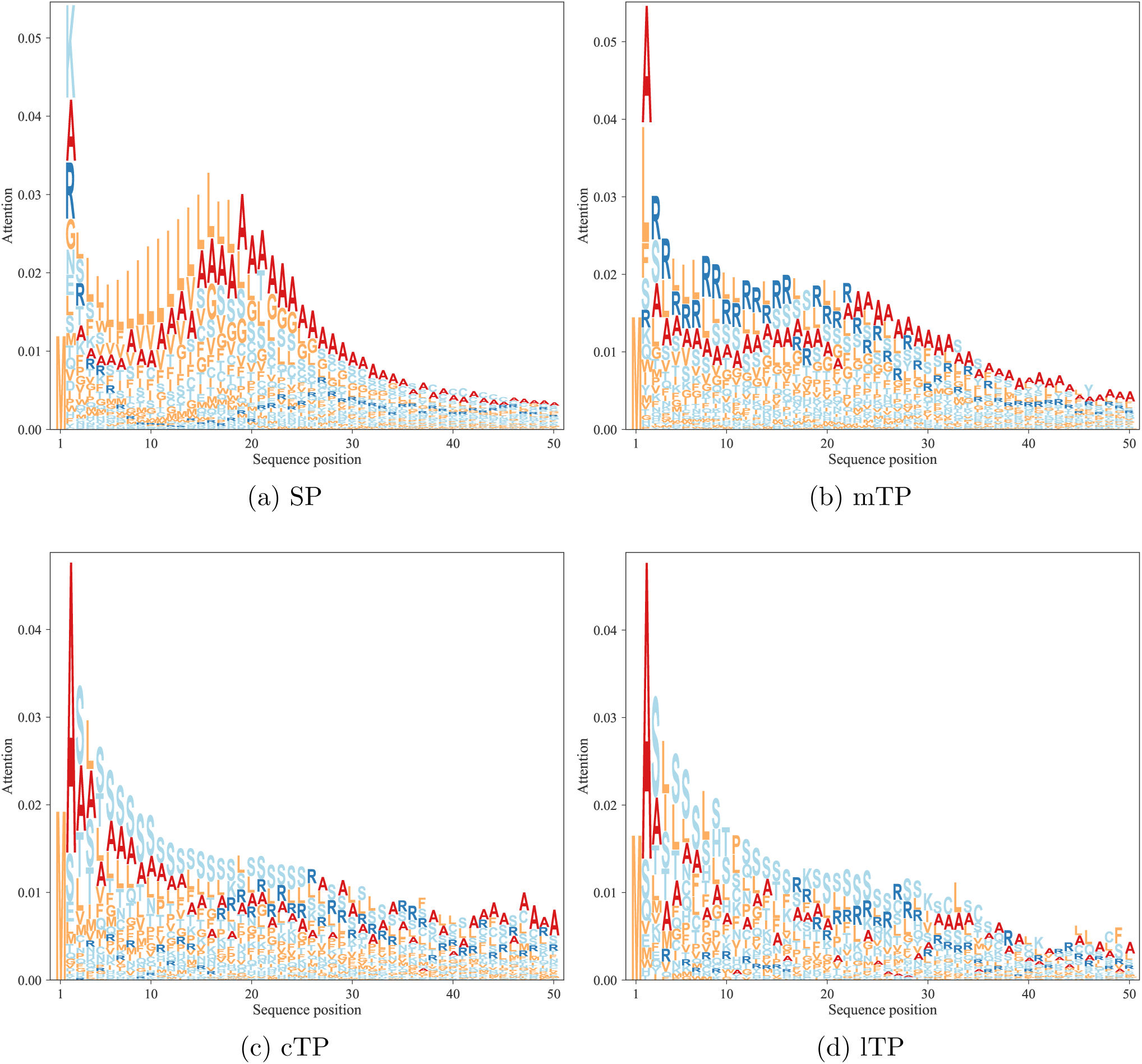
Attention layer LOGOs showing the impact strength of the attention layer and the frequency of amino acids. All sequences are aligned at the N-terminus.

### 3.5. The R-3 rule appears important for mTPs

In Figure 5 it can be seen that the attention layer focuses on the position just before the cleavage site (the −1 position). In SPs, cTPs and lTPs position −1 is dominated by alanine, while in mTPs this position is dominated by tyrosine and phenylalanine. In contrast, the actual cleavage signal is dominated by a couple of positions (such as −1 and −3 in SPs and lTPs), see Figure 7 and S2. This difference can be explained by the attention layer collapsing information from nearby positions into one position. In addition to the site close to the cleavage site, most of the information obtained from the attention layers is directly N-terminal of the cleavage site. In agreement with what is known about the differences between the targeting peptides, the attention for the SPs is focused on a stretch of approximately 10 hydrophobic residues, while the other peptides have a longer stretch of informative residues. As is well known, the mTPs are enriched in arginine.

**Figure 7:**
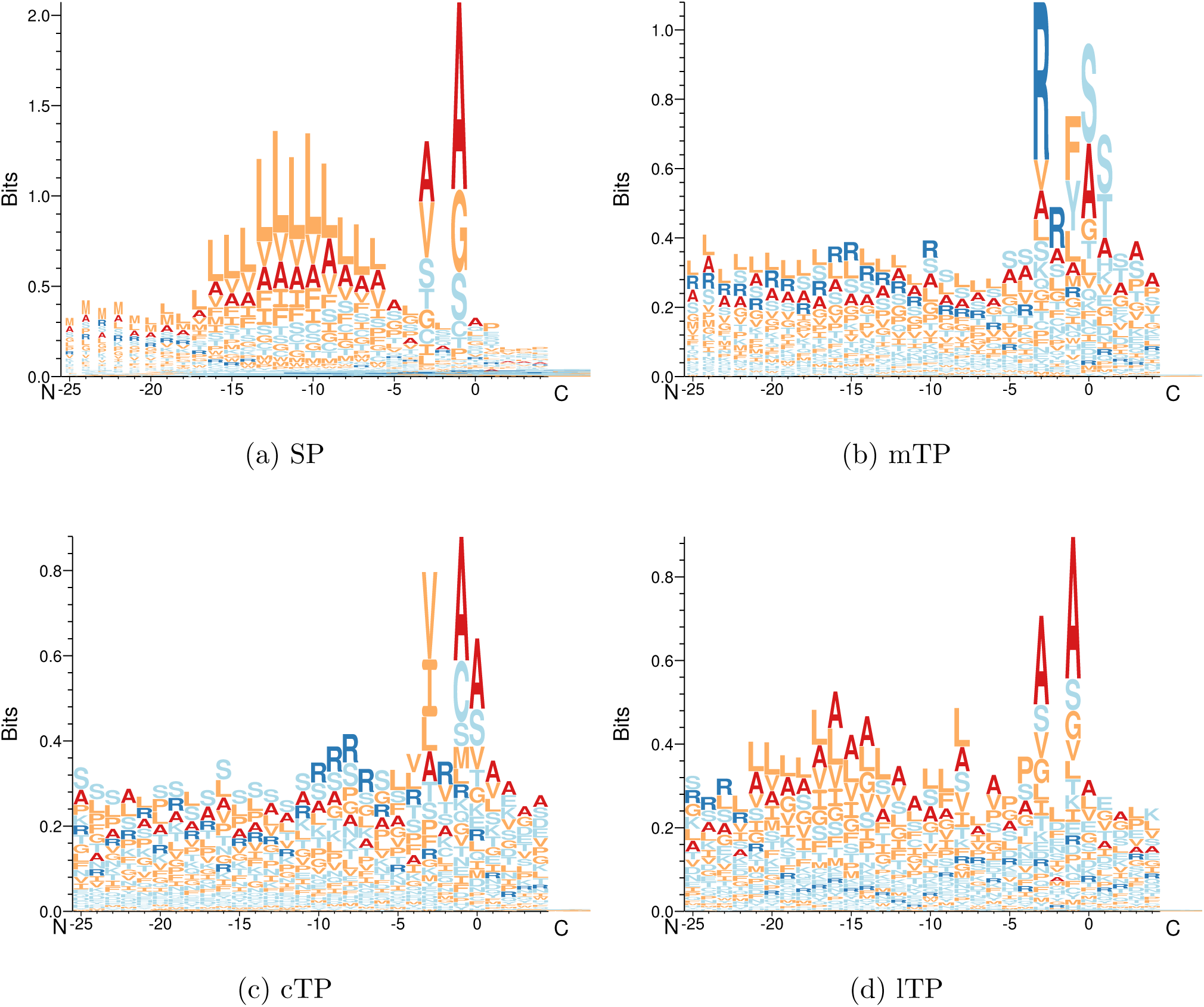
Sequence LOGOs showing the amino acid frequencies in the pre-sequences. All sequences are aligned according to the predicted cleavage site.

### 3.6. TargetP 2.0 overpredicts mTP cleavage sites with arginine in −3

For the SPs, cTPs and lTPs, the sequence logos are almost identical between predicted and experimentally annotated proteins, both in the cleavage site and the signal composition. However, we can observe that for mTPs the amino acid composition near the cleavage site differs between predicted, Figure 7b, and experimentally verified mTPs, Figure S2b. In both cases, there is an abundance of arginines in position −2, −3 and −10 from the cleavage site as described before [31, 32]. However, the signal for arginine at −3 is stronger among the predicted than among the experimentally verified cleavage sites. In order to investigate this difference further, we plotted the distribution of the distance from the experimental and predicted cleavage sites to the nearest upstream arginine, see Figure S3. It shows that while there is good agreement at most positions, there is a clear over-prediction at −3 and an under-prediction at −10.

The sites with arginine at −2 are thought to represent the original cleavage by Mitochondrial Processing Peptidase (MPP), while the sites with arginine at −3 and −10 are thought to arise by subsequent cleavage events by the Icp55 peptidase and Mitochondrial Intermediate Peptidase (MIP), respectively [33, 32, 34, 35, 8]. The cleavage by Icp55 could explain the fact that some patterns in the mTP cleavage site, Figures 7b and S2b, seem to be repeated with a shift of one position, e.g. the preference for serine that occurs in positions one and two in the mature protein.

The findings represented in Figure S3 show that the model can easily recognise the arginines at position −2 (original MPP sites) and −3 (Icp55 sites) but has troubles in identifying arginines at position −10 (MIP sites). This over-representation of arginine at position −3 and under-representation at position −10 is probably contributing to the relatively low performance on the cleavage site prediction in mTPs. It might be relevant to explore further the distance of arginines from the cleavage site and the patterns recognised by the three peptidases to improve the prediction of the mTP cleavage site in future versions.

### 3.7. cTPs have an alanine in position two

There is also a strong attention peak at position two for all targeting peptides, see Figure 6. From the sequence logo it is clear that position two amino acid preferences differ between targeting peptides, see Figures 8 and S4. In cTPs and lTPs there is a powerful signal for alanine in position two. In contrast, signal peptides have some preference for lysine and mTPs for alanine or leucine in position two. see Table S8.

**Figure 8:**
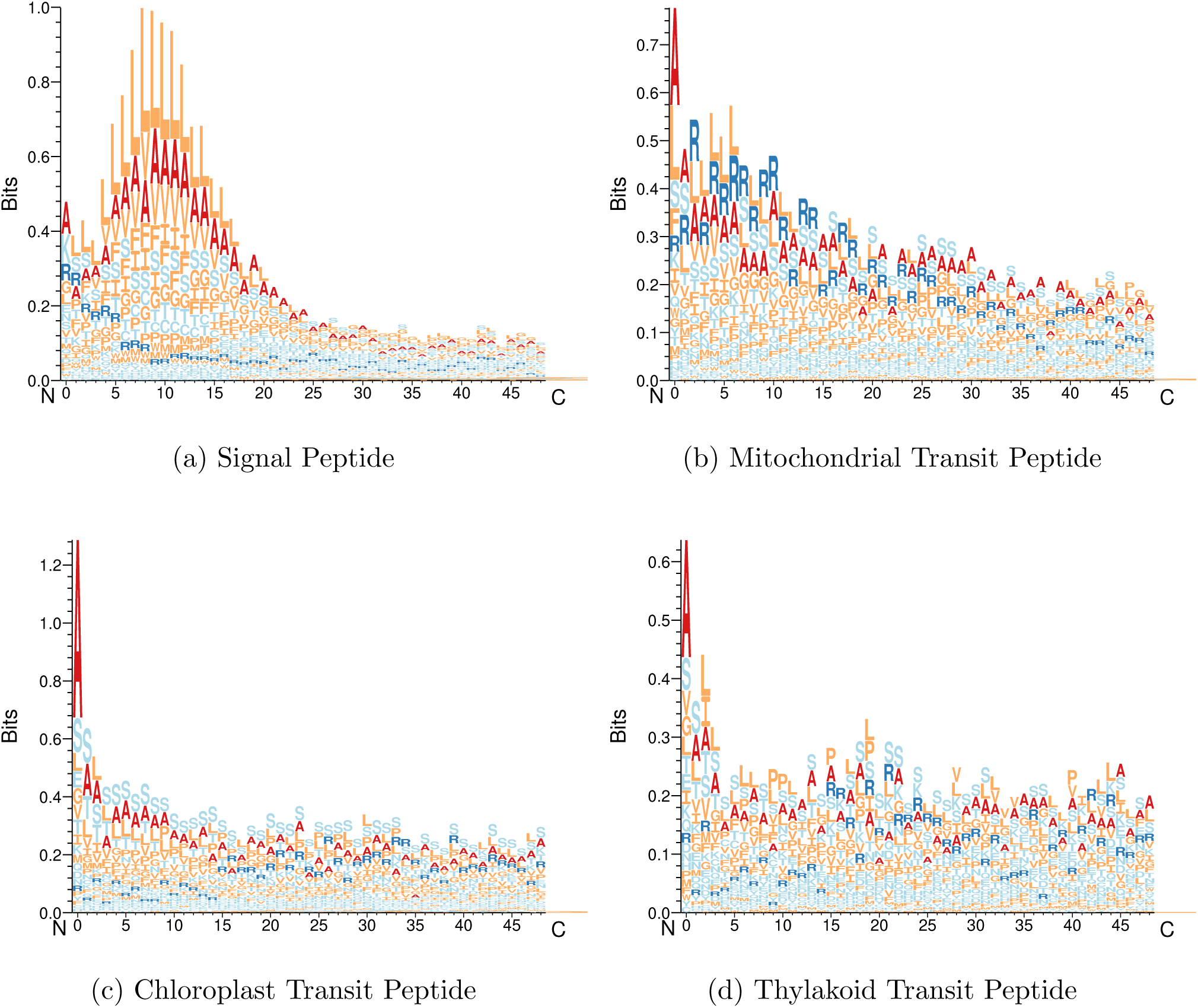
Sequence LOGOs showing the amino-terminal pre-sequences. All sequences are aligned at the N-terminus.

The importance of position two is likely to be related to the cleavage of the N-terminal methionine. When there is a short side-chained amino acid (Ala, Cys, Gly, Pro, or Ser) in position two, the methionine can be cleaved by a methionine aminopeptidase (MAP) [36]. There exist two classes of MAPs, MAP1 and MAP2. All these proteins are homologous to the machinery in bacteria, indicating that they work co-translationally. *A. thaliana* has four MAP1s (MAP1A, MAP1B, MAP1C and MAP1D) and two MAP2s (MAP2A and MAP2B). It has been shown that MAP1B, MAP1C and MAP1D are targeted for proteins belonging to the organelles [37].

In Figure 1 it can be seen that about 60% of the proteins without targeting peptides have an amino acid in position two that allows the N-methionine to be cleaved. These proteins have mostly alanine or serine in position two. The N-terminal methionine can only be cleaved if the second residue has a short side-chain. For proteins with signal peptides, in all species except the plants, less than 40% of the residues in position two have a short side-chain. The same can be seen for mTPs in the fungi single-celled eukaryotic groups. Most striking is the observation that about two thirds of the cTPs and lTPs have an alanine in position two, see Figure 1. This preference has been noted before [38, 39]. When mutating the second position in dual-targeting proteins that are imported to both chloroplasts and mitochondria the targeting was disrupted [40]. Surprisingly, when the authors mutated one of the few chloroplast proteins that did not have an alanine in position two, *PheRS*, from threonine to alanine the import to chloroplasts decreased.

It has been reported that amino acid frequencies in position two differ between species [41]. The frequency of alanine in position two varies from 7% in *Escherichia coli* to close to 30% in *A. thaliana*. In table S9 it can be seen that alanine is frequent in all types of proteins in *A. thaliana* but also that the frequency is higher in proteins targeted for plastids. One possible reason for alanine to be preferred in position two is that alanine has a strong helical propensity. The amino-terminal sections of cTPs and lTPs are less prone to form secondary structures than mTPs and SPs at the amino-terminal, see Figure S5. Here, it can also be seen that signal peptides have a much stronger tendence to form structure close to the N-terminal than the other peptides. The importance of the N-termini can also be seen by the fact that the simple MLP-20 method performs quite well at identification of noTPs, SPs and mTPs. However, to fully understand the importance of the second position additional experimental studies are needed.

## 4. Conclusions

Here, we introduce the new version of TargetP 2.0 that includes the prediction of thylakoid transit peptides and uses deep neural networks. TargetP 2.0 can be helpful to accurately annotate N-terminal sorting signals and cleavage sites in particular as it scales to complete proteomes. TargetP 2.0 outperforms all other methods in all N-terminal sorting signals. Regarding classification the only alternative method that comes close to TargetP 2.0 in performance is DeepLoc for signal peptides and mitochondrial targeting peptides. However, for chloroplast peptides, TargetP 2.0 is superior, and DeepLoc does not predict thylakoid localisation. On the other hand, DeepLoc also predicts many other subcellular localisations not governed by targeting peptides.

When analysing how TargetP 2.0 arrives at its predictions, we note that two distinct regions contribute. As expected, the region around the cleavage site is essential for classification of the type of transit peptide. However, surprisingly, an equally important contribution comes from the N-terminal region. Upon closer inspections, it is clear that (i) in plants, two-thirds of the chloroplast and thylakoid targeting peptides have an alanine in position two (after the N-terminal methionine), and (ii) in fungi only 20-30% of the N-termini of mTPs and SPs can be cleaved, compared to 60% for proteins without targeting peptides. In summary, this indicates that it is not unlikely that specificity of methionine aminopeptidases aids in the co-translational targeting of peptides into organelles.

## 5. Acknowledgments

We thank Castrense Savojardo for having run TPPred3 for our analysis. We thank Elzbieta Glaser for discussions about earlier work regarding the importance of alanine in position 2 in cTPs. We thank the NVIDIA Corporation and the Swedish National Infrastructure for Computing for providing computational resources. AE was supported by grant VR-NT-2016-03798 from the Swedish National Research Council.

## 8. Supplementary Figures

**Figure S1:**
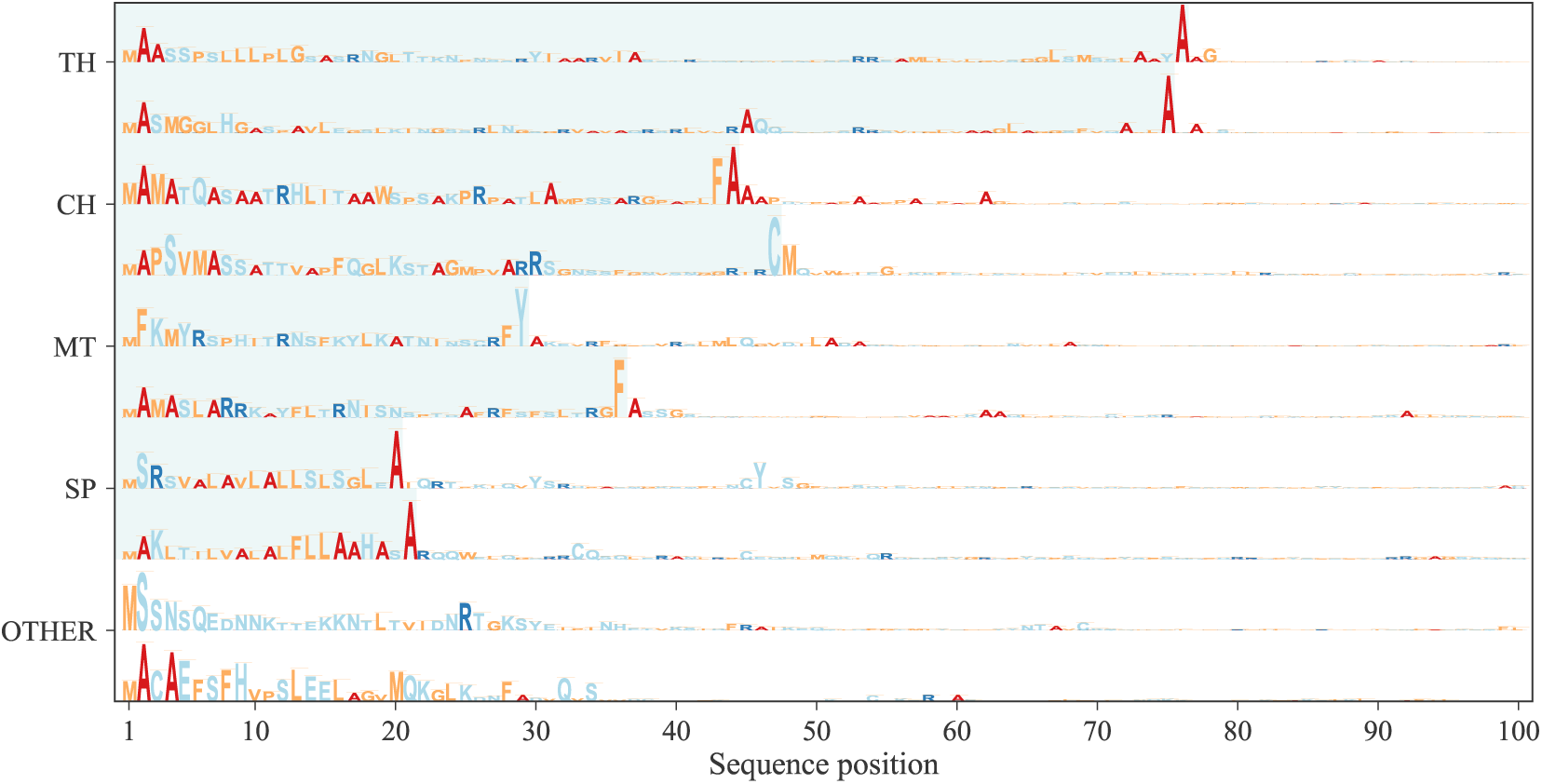
Representation of the attention weights for a few proteins. The height of the letter represents the attention weight in that position and the letter the type of amino acid. The shaded area corresponds to the predicted targeting peptide (SP, mTP, cTP or lTP).

**Figure S2:**
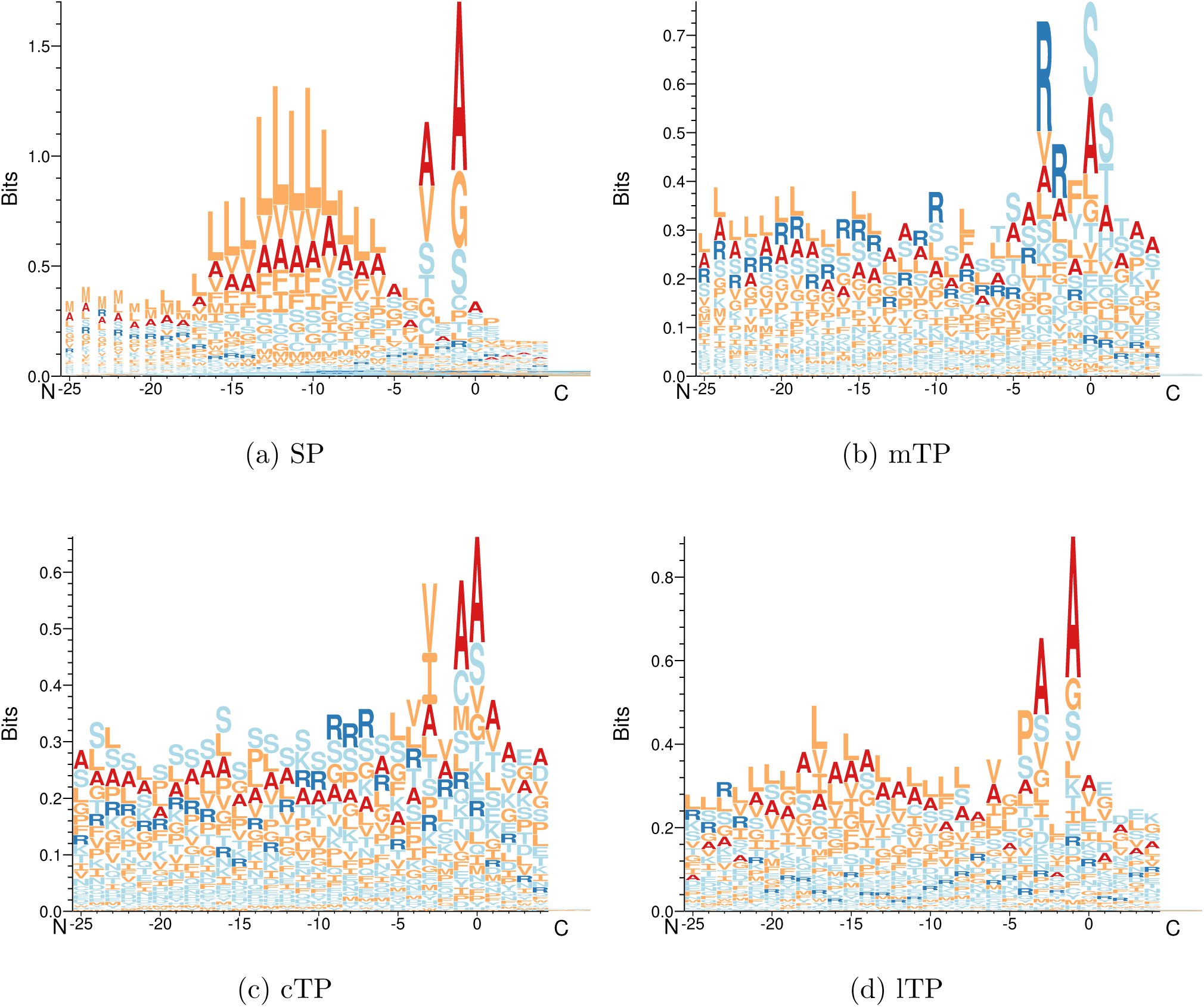
Sequence LOGOs showing the experimental amino-terminal pre-sequences. Sequences are aligned according to the annotated cleavage site.

**Figure S3:**
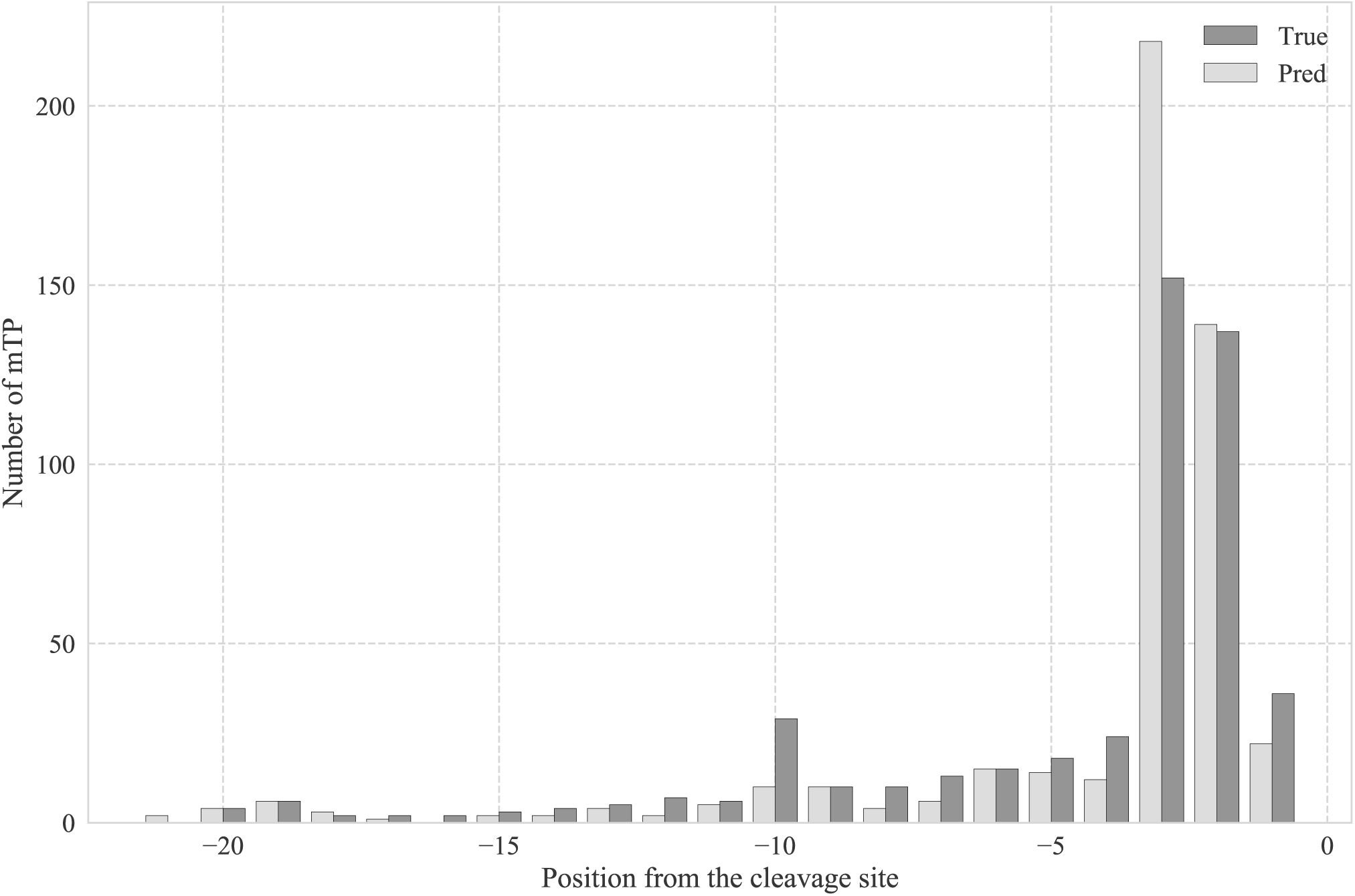
Distribution of the distance from true and predicted cleavage sites to the nearest arginine in mitochondrial transit peptides.

**Figure S4:**
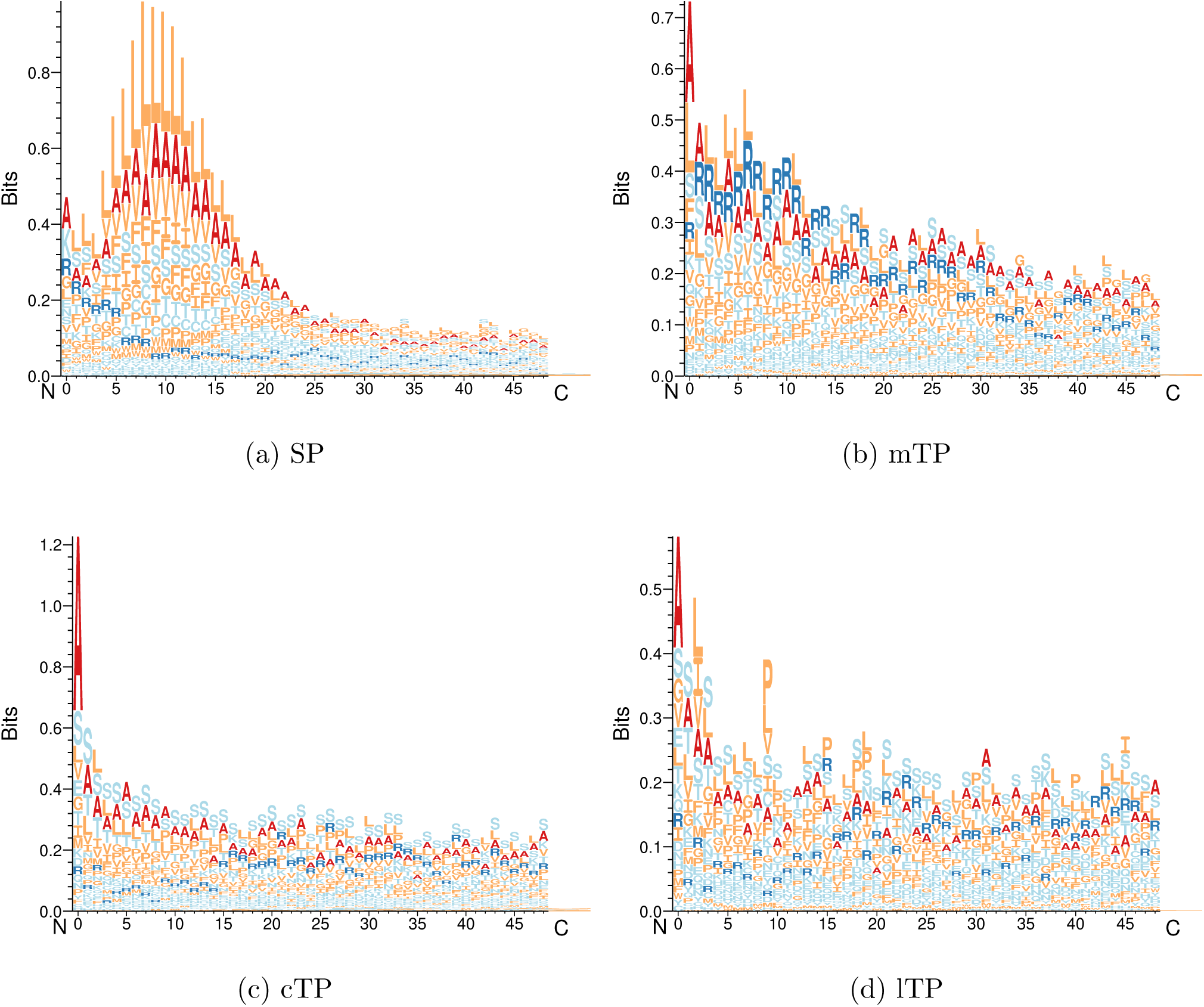
Sequence LOGOs showing the experimental amino-terminal pre-sequences. Sequences are aligned at the N-terminus.

**Figure S5:**
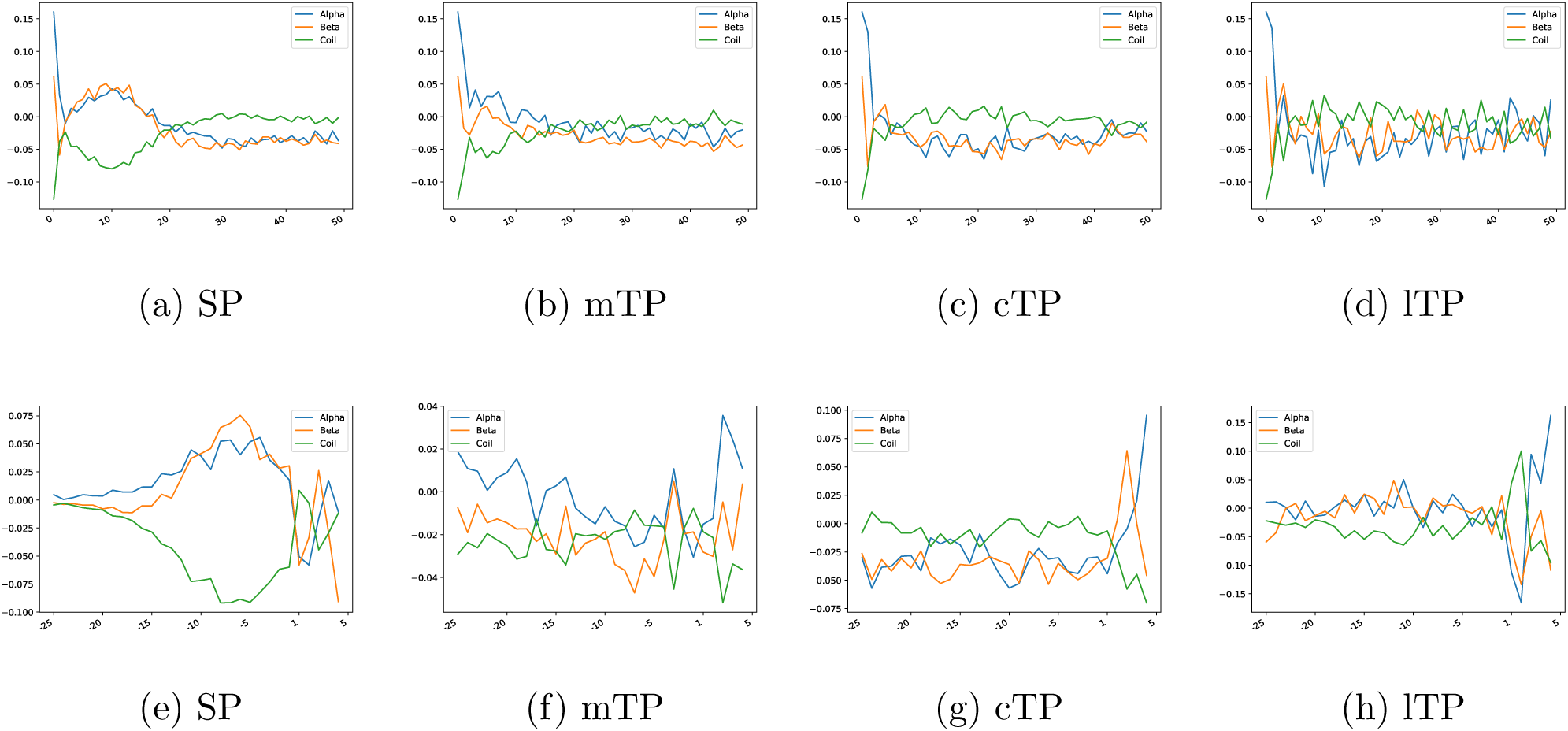
Log-odds ratio of secondary structure preferences for the different targeting peptides. Upper row shows the peptides aligned at N-terminal and the lower row show the peptides aligned at the cleavage site.

## 9. Supplementary Tables

**Table S1:** F1 score for the MLP predictor, using different number of N-terminal residues.

|  | **Tool** | **SP** | **mTP** | **cTP** | **lTP** | **noTP** | **Average** |
| --- | --- | --- | --- | --- | --- | --- | --- |
|  | **MLP-5** | 0.57 | 0.22 | 0.03 | 0.00 | 0.88 | 0.77 |
|  | **MLP-10** | 0.80 | 0.55 | 0.33 | 0.04 | 0.93 | 0.87 |
|  | **MLP-15** | 0.90 | 0.57 | 0.42 | 0.00 | 0.95 | 0.91 |
|  | **MLP-20** | 0.93 | 0.63 | 0.43 | 0.04 | 0.96 | 0.93 |

**Table S2:** Confusion matrix for non-plant organisms representing the number of proteins for each targeting peptides predicted by TargetP 2.0 (rows) versus observed in the test set (columns).

|  | **Class** | **SP** | **mTP** | **noTP** |
| --- | --- | --- | --- | --- |
|  | **SP** | 2390 | 5 | 47 |
|  | **mTP** | 2 | 311 | 44 |
|  | **noTP** | 23 | 58 | 7644 |

**Table S3:**
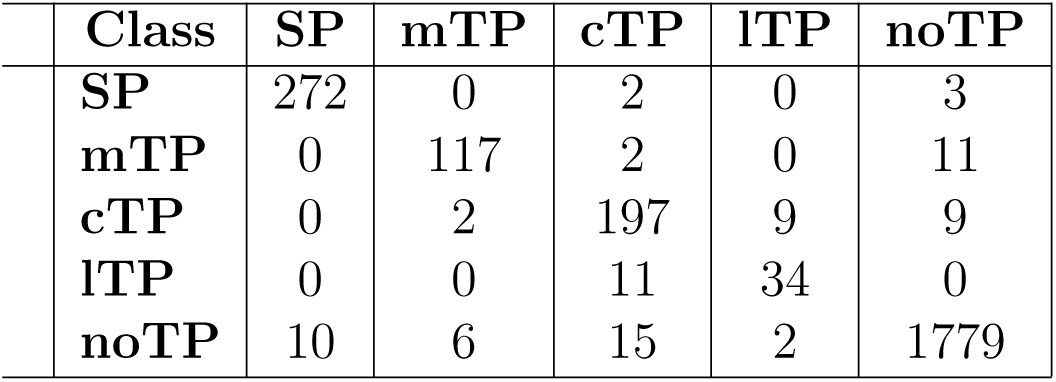
Confusion matrix for Viridiplantae representing the number of proteins for each targeting peptides predicted by TargetP 2.0 (rows) versus observed in the test set (columns).

|  | **Class** | **SP** | **mTP** | **cTP** | **lTP** | **noTP** |
| --- | --- | --- | --- | --- | --- | --- |
|  | **SP** | 272 | 0 | 2 | 0 | 3 |
|  | **mTP** | 0 | 117 | 2 | 0 | 11 |
|  | **cTP** | 0 | 2 | 197 | 9 | 9 |
|  | **lTP** | 0 | 0 | 11 | 34 | 0 |
|  | **noTP** | 10 | 6 | 15 | 2 | 1779 |

**Table S4:** Performance of TargetP 2.0 considering only the peptide prediction in one kingdom at a time. The table shows the performance in the test set yield by each predictor for Mitochondria (mTP), Chloroplast (cTP), Thylakoid (lTP), Signal Peptide (SP), and the other proteins without targeting peptide (noTP), in terms of F1 score, Matthew correlation coefficient (Mcc), Precision (Prec) and Recall (Rec)

|  | **Kingdom** | **Loc** | **Proteins** | **Prec** | **Rec** | **F1-Score** | **MCC** |
| --- | --- | --- | --- | --- | --- | --- | --- |
|  | **Viridiplantae** | SP | 282 | 0.98 | 0.96 | 0.97 | 0.97 |
|  | **Metazoa** | SP | 2251 | 0.98 | 0.99 | 0.99 | 0.98 |
|  | **Fungi** | SP | 133 | 0.91 | 0.99 | 0.95 | 0.95 |
|  | **Other** | SP | 31 | 1.0 | 0.97 | 0.98 | 0.98 |
|  | **Viridiplantae** | mTP | 125 | 0.9 | 0.94 | 0.92 | 0.91 |
|  | **Metazoa** | mTP | 263 | 0.89 | 0.86 | 0.87 | 0.87 |
|  | **Fungi** | mTP | 103 | 0.83 | 0.77 | 0.80 | 0.79 |
|  | **Other** | mTP | 8 | 0.88 | 0.88 | 0.88 | 0.87 |
|  | **Viridiplantae** | cTP | 227 | 0.91 | 0.87 | 0.89 | 0.88 |
|  | **Viridiplantae** | lTP | 45 | 0.76 | 0.76 | 0.76 | 0.75 |
|  | **Viridiplantae** | noTP | 1802 | 0.98 | 0.99 | 0.98 | 0.94 |
|  | **Metazoa** | noTP | 5354 | 0.99 | 0.99 | 0.99 | 0.97 |
|  | **Fungi** | noTP | 2263 | 0.99 | 0.99 | 0.99 | 0.88 |
|  | **Other** | noTP | 118 | 0.98 | 0.99 | 0.99 | 0.95 |

**Table S5:** Performance of the predictors considering the peptide and cleavage site. The table shows the performance in the test set yield by each predictor for Mitochondria (mTP), Chloroplast (cTP), Thylakoid (lTP) and Signal Peptide(SP) in terms of Precision (Prec) and Recall (Rec) both for the cleavage site (CS) and targeting peptide (PEP).

|  | Tool | Loc | Recall | No. Correct | No. Correct -5/+5 | Recall -5/+5 | Total No. |
| --- | --- | --- | --- | --- | --- | --- | --- |
|  | **TargetP 2.0** | **SP** | 0.83 | 2248 | 2584 | 0.96 | 2697 |
|  | **TargetP 1.1** | **SP** | 0.83 | 2250 | 2551 | 0.95 | 2697 |
|  | **PredSL** | **SP** | 0.70 | 1889 | 2312 | 0.86 | 2697 |
|  | **TargetP 2.0** | **mTP** | 0.46 | 230 | 326 | 0.65 | 499 |
|  | **TargetP 1.1** | **mTP** | 0.42 | 211 | 276 | 0.55 | 499 |
|  | **PredSL** | **mTP** | 0.16 | 81 | 196 | 0.39 | 499 |
|  | **TPPred3** | **mTP** | 0.39 | 195 | 259 | 0.52 | 499 |
|  | **Mitofates** | **mTP** | 0.18 | 88 | 251 | 0.50 | 499 |
|  | **TargetP 2.0** | **cTP** | 0.49 | 111 | 164 | 0.72 | 227 |
|  | **TargetP 1.1** | **cTP** | 0.07 | 17 | 104 | 0.46 | 227 |
|  | **PredSL** | **cTP** | 0.11 | 25 | 67 | 0.30 | 227 |
|  | **TPPred3** | **cTP** | 0.30 | 67 | 105 | 0.46 | 227 |
|  | **TargetP 2.0** | **lTP** | 0.60 | 27 | 31 | 0.69 | 45 |
|  | **PredSL** | **lTP** | 0.10 | 5 | 32 | 0.71 | 45 |

**Table S6:**
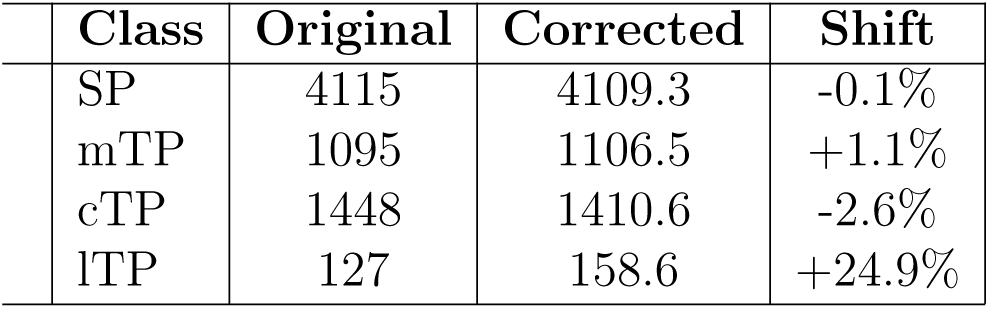
Corrected number of proteins annotated with different targeting peptides for the A. thaliana genome using the confusion matrix from Table S3.

**Table S7:** The table shows the the agreement with Uniprot annotations.

|  | Kingdom | Organism | Reference | Peptide | TargetP 2.0 | Uniprot | Agree |
| --- | --- | --- | --- | --- | --- | --- | --- |
|  | Metazoa | HUMAN | 20585 | SP | 3698 | 3521 | 3382 |
|  | Metazoa | DROME | 13785 | SP | 3323 | 3076 | 2994 |
|  | Metazoa | MOUSE | 22286 | SP | 4278 | 4042 | 3883 |
|  | Metazoa | CAEEL | 19986 | SP | 4591 | 4078 | 3967 |
|  | Metazoa | XENTR | 24138 | SP | 2366 | 1933 | 1841 |
|  | Metazoa | DANRE | 25747 | SP | 4399 | 3808 | 3651 |
|  | Fungi | YEAST | 6049 | SP | 386 | 298 | 272 |
|  | Fungi | SCHPO | 5142 | SP | 252 | 214 | 195 |
|  | Viridiplantae | ARATH | 27623 | SP | 4115 | 3543 | 3374 |
|  | Viridiplantae | BRADI | 34230 | SP | 3987 | 3567 | 3216 |
|  | Viridiplantae | ORYSJ | 43588 | SP | 4687 | 4169 | 3644 |
|  | Viridiplantae | SOLLC | 33952 | SP | 3904 | 2848 | 2675 |
|  | Viridiplantae | VITVI | 29882 | SP | 2980 | 2199 | 2019 |
|  | Metazoa | HUMAN | 20585 | mTP | 627 | 540 | 442 |
|  | Metazoa | DROME | 13785 | mTP | 522 | 136 | 102 |
|  | Metazoa | MOUSE | 22286 | mTP | 631 | 519 | 429 |
|  | Metazoa | CAEEL | 19986 | mTP | 447 | 116 | 88 |
|  | Metazoa | XENTR | 24138 | mTP | 453 | 51 | 37 |
|  | Metazoa | DANRE | 25747 | mTP | 626 | 98 | 70 |
|  | Fungi | YEAST | 6049 | mTP | 368 | 365 | 284 |
|  | Fungi | SCHPO | 5142 | mTP | 250 | 266 | 159 |
|  | Viridiplantae | ARATH | 27623 | mTP | 1095 | 526 | 432 |
|  | Viridiplantae | BRADI | 34230 | mTP | 970 | 0 | 0 |
|  | Viridiplantae | ORYSJ | 43588 | mTP | 1100 | 86 | 67 |
|  | Viridiplantae | SOLLC | 33952 | mTP | 931 | 4 | 4 |
|  | Viridiplantae | VITVI | 29882 | mTP | 725 | 0 | 0 |
|  | Viridiplantae | ARATH | 27623 | cTP | 1448 | 1222 | 884 |
|  | Viridiplantae | BRADI | 34230 | cTP | 1781 | 0 | 0 |
|  | Viridiplantae | ORYSJ | 43588 | cTP | 2049 | 340 | 279 |
|  | Viridiplantae | SOLLC | 33952 | cTP | 1274 | 78 | 57 |
|  | Viridiplantae | VITVI | 29882 | cTP | 1125 | 3 | 2 |
|  | Viridiplantae | ARATH | 27623 | ITP | 127 | 72 | 58 |
|  | Viridiplantae | BRADI | 34230 | ITP | 85 | 0 | 0 |
|  | Viridiplantae | ORYSJ | 43588 | ITP | 84 | 9 | 5 |
|  | Viridiplantae | SOLLC | 33952 | ITP | 117 | 1 | 1 |
|  | Viridiplantae | VITVI | 29882 | ITP | 91 | 0 | 0 |

**Table S8:**
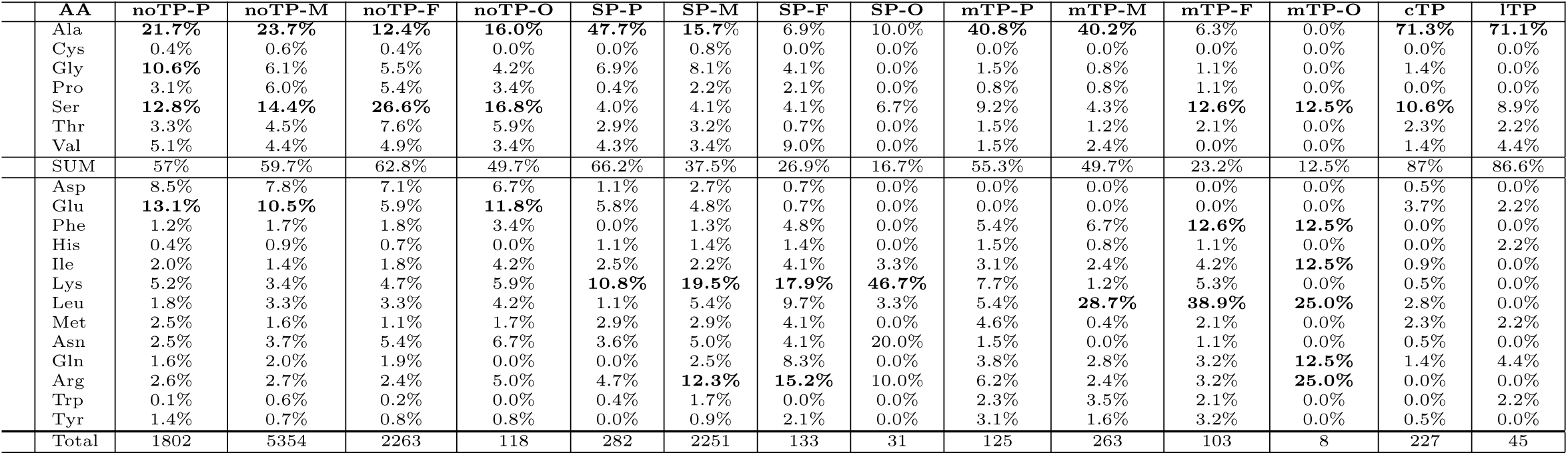
Frequencies in position two in the test set divided by Viridiplantae (P), Metazoa (M), Fungi (F) and other Eukaryotic organisms (O) sequences for the five categories of proteins, mitochondrial Transit Peptides (mTPs), Signal Peptide (SP), choroplast Transit Peptides (cTP), thylakoid lumenal Tranist Peptides (lTP) and proteins with no targeting peptide (noTP). All frequencies higher than 10% are marked in bold. The top part shows the frequency of the short-chained amino acids that can be cleaved by MAPs. The SUM line is the sum of all these short-chained amino acids, and the Total line is the number of proteins in each class.

**Table S9:** Predicted frequencies in position two in A. thaliana by TargetP 2.0. All frequencies higher than 10% are marked in bold.

|  | AA | noTP | SP | mTP | cTP | lTP |
| --- | --- | --- | --- | --- | --- | --- |
|  | A | **16.4%** | **25.7%** | **27.4%** | **48.3%** | **32.1%** |
|  | C | 1.0% | 0.7% | 0.4% | 0.7% | 3.6% |
|  | D | 8.1% | 3.8% | 0.0% | 1.0% | 3.6% |
|  | E | **13.0%** | 8.5% | 0.8% | 5.0% | **10.7%** |
|  | F | 2.1% | 2.6% | 6.6% | 0.7% | 3.6% |
|  | G | **11.0%** | 7.6% | 1.9% | 2.0% | 0.0 % |
|  | H | 0.9% | 0.7% | 0.8% | % | 3.6% |
|  | I | 2.6% | 2.7% | 3.1% | 2.7% | 3.6% |
|  | K | 5.2% | **11.1%** | 3.5% | 1.3% | 3.6% |
|  | L | 3.5% | 2.9% | **11.6%** | 6.0% | 3.6% |
|  | M | 2.4% | 2.5% | 3.5% | 2.0% | 3.6% |
|  | N | 3.5% | 3.4% | 3.5% | 1.3% | 3.6% |
|  | P | 2.6% | 1.0% | 0.4% | 1.3% | 3.6% |
|  | Q | 1.8% | 1.1% | 3.1% | 1.7% | 3.6% |
|  | R | 2.9% | 2.9% | 7.7% | 0.3% | 0.0 % |
|  | S | **10.1%** | 9.3% | **15.4%** | **14.8%** | 3.6% |
|  | T | 4.4% | 5.8% | 2.7% | 4.0% | 3.6% |
|  | V | 6.2% | 6.2% | 2.3% | 4.4% | 7.1% |
|  | W | 0.8% | 0.5% | 1.9% | 1.0% | 0.0 % |
|  | Y | 1.4% | 0.8% | 3.5% | 1.3% | 3.6% |

## Abbreviations

SP: Signal Peptide
mTP: mitochondrial Transit Peptide
cTP: chloroplast Transit Peptide
lTP: thylakoid lumenal Transit Peptide
noTP: proteins without Targeting Peptide
RNN: Recurrent Neural Network
LSTM: Long Short-Term Memory

